# SPECS: A non-parameteric method to identify tissue-specific molecular features for unbalanced sample groups

**DOI:** 10.1101/656397

**Authors:** Celine Everaert, Pieter-Jan Volders, Annelien Morlion, Olivier Thas, Pieter Mestdagh

## Abstract

To understand biology and differences among various tissues or cell types, one typically searches for molecular features that display characteristic abundance patterns. Several specificity metrics have been introduced to identify tissue-specific molecular features, but these either require an equal number of replicates per tissue or they can’t handle replicates at all. We describe a non-parametric specificity score that is compatible with unequal sample group sizes. To demonstrate its usefulness, the specificity score was calculated on all GTEx samples, detecting known and novel tissue-specific genes. A webtool was developed to browse these results for genes or tissues of interest. An example python implementation of SPECS is available at https://github.ugent.be/ceeverae/SPECs. The precalculated SPECS results on the GTEx data are available through a user-friendly browser at specs.cmgg.be.

## 1 Introduction

To understand biology and differences among various tissues or cell types, one typically searches for molecular features (i.e. RNA, protein, metabolites) that display characteristic abundance patterns. In the most extreme case, these features display tissue-or cell-type restricted abundance profiles. Such specific features can provide insights in functional, development or disease mechanisms (Leucci *et al.*, 2016) or serve as biomarkers (Stutterheim *et al.*, 2008; Prensner *et al.*, 2013). Various consortium-based efforts have generated vast amounts of molecular data that can be exploited for this purpose. The Genotype-Tissue Expression (GTEx) project (https://gtexportal.org) and The Cancer Genome Atlas (TCGA) (https://www.cancer.gov/tcga) are examples of such rich resources containing RNA-sequencing based molecular features for thousands of samples derived from various individuals and tissue types (Lonsdale *et al.*, 2013). To identify tissue-specific molecular features, several specificity metrics have been introduced, but these can suffer from data loss introduced by the requirement to collapse data from biological replicates. Moreover, those metrics that can handle biological replicates require equal sample sizes. In this application note, we describe a novel non-parametric specificity score that is compatible with unequal sample group sizes and enables the detection of features that are specifically present or absent in one or more tissue types.

## 2 Methods

Let the index *d* = 1,…,m_d_ refer to a particular sample state. Depending on the application and whether the user wants to give weight to a certain state, π_d_ is the prevalence of state *d* in the target population or π_d_ is equilibrated. Suppose there are m_g_ candidate features, i.e. g = 1,…,m_g_. Let Y_gd_ denote the outcome of feature *g* in state *d* with n_gd_ observations, so that the individual outcomes are denoted by Y_gdi_, i = 1, …,n_gd_. The Y_g-d_ notation denotes the outcome of feature *f* in all groups but the state *d*. The index *g* will be dropped in further notations. A feature is a characteristic for a given state if its outcome distribution for the given state shows no overlap with the outcome distributions of the other states. This means a larger AUC, given by:

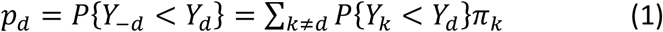

If p_d_ is close to zero or one, the distributions are well separated. The probabilities P{Y_k_<Y_d_} are computationally fast to calculate. The probability P_kd_ = P{Y_k_<Y_d_} is then estimated as:

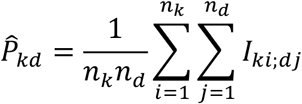

with *I*_ki;dj_ a 0/1 indicator for the event *Y*_*ki*_<*Y*_*dj*_.

Hence, an estimator of p_d_ is given by:

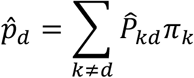

Further selection of features can be performed based on the distributions of 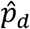 as explained in Supplemental Methods 1. As this is a computationally intensive step for large data matrices, one can opt to select features based on a threshold. In our use case, we defined state-specific features as those where the score 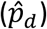 for one state was above 0.95 and features that were specifically absent in one state as those with a score 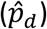 lower than 0.05. If the score of 0.95 or 0.05 was reached in multiple states, the feature was defined as specific (present or absent) for all these states. The python implementation of the method is available at https://github.ugent.be/ceeverae/SPECs.

## 3 Results

The Genotype-Tissue Expression (GTEx v7) project (Lonsdale *et al.*, 2013) consists of RNA sequencing data from 12 766 samples belonging to 31 different tissues (7 to 1854 samples per tissue). We calculated the SPECS specificity score on normalized counts for all Ensembl (GRCh38.v85) genes (n=56 202) using all samples. For 30 of the 31 tissues, 2 (esophagus) to 7948 (testis) specifically expressed genes were identified. Most of these genes are protein coding (n=10 959), followed by lincRNAs (n=3080), antisense genes (n=2022) and pseudogenes (n=1976) (Figure 1A and Supplemental Figure 1). In addition, the method has the ability to identify genes that are highly specific for two (or more) tissues, with specificity scores that are slightly lower. As expected, the tissues with the highest number of common specific genes are biologically related such as spleen and blood, or brain and pituitary or muscle and hart.

**Figure 1.**
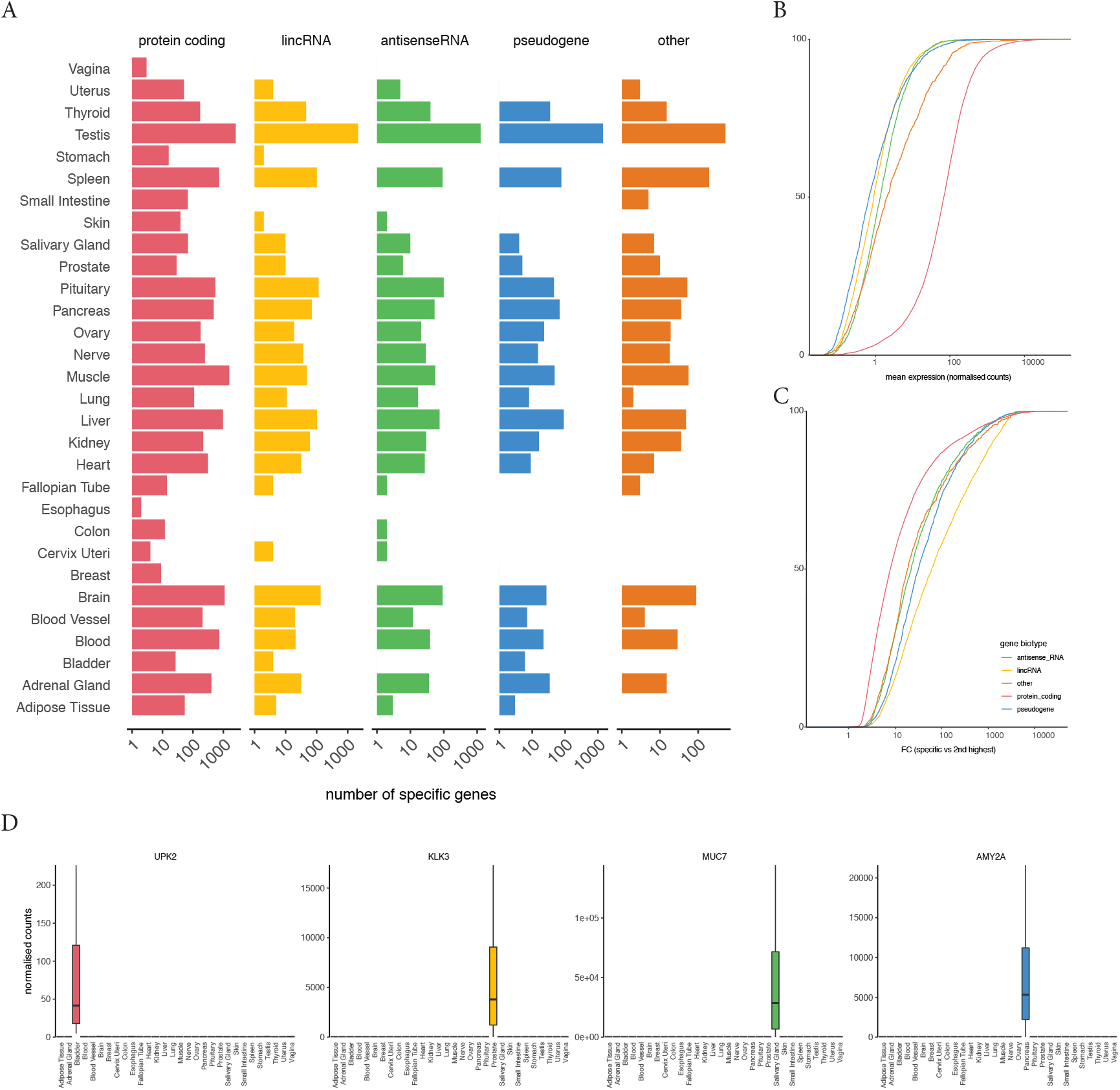
Known and novel genes are detected as specific for various biotypes. A) The number of specific genes for each GTEx tissue and biotype shows that most specific genes are protein-coding. B) Cumulative distribution of the mean expression of specific genes, shows that specific protein-coding genes are higher expressed compared to the other biotypes. C) Cumulative distribution of the fold changes of specific genes and the 2nd tissue shows larger differences for lincRNA genes compared to other biotypes. D) Examples of well-known specific genes; UPK2 for bladder, KLK3 for prostate, MUC7 for adrenal gland and AMY2A for pancreas.

Besides genes that are specifically abundant in a tissue, our method also enables the identification of genes that are specifically repressed in a given tissue. These so-called disallowance genes (Thorrez *et al.*, 2010) were found for 17 tissues ranging from 2 (salivary gland) to 1989 (blood) genes. Most of these are protein coding genes (Supplemental Figure 2).

For all specifically abundant genes we calculated fold changes between the specific tissue(s) and all other tissues. The fold changes for lincRNAs were typically higher than for other biotypes, in line with previous studies in which lincRNAs were shown to be more specific compared to protein coding genes (Cabili *et al.*, 2011) (Figure 1B and Figure 1C). From our analyses, known specific genes are readily confirmed, such as kallikrein related peptidase 2 (KLK2) and 3 (KLK3, also known as PSA) for prostate, uroplakin 2 (UPK2) for bladder, mucin 7 (MUC7) for the salivary gland and amylase alpha 2A (AMY2A) for pancreas (Figure 1D). For each tissue in GTEx, rank percentiles for the specific genes are pre-calculated and distilled into a web tool (specs.cmgg.be) where a user can select either their gene of interest to evaluate its specificity or a tissue of interest to identify the most specific genes.

