## Supplemental Figures for "SPECS: A non-parameteric method to identify tissue-specific molecular features for unbalanced sample groups"

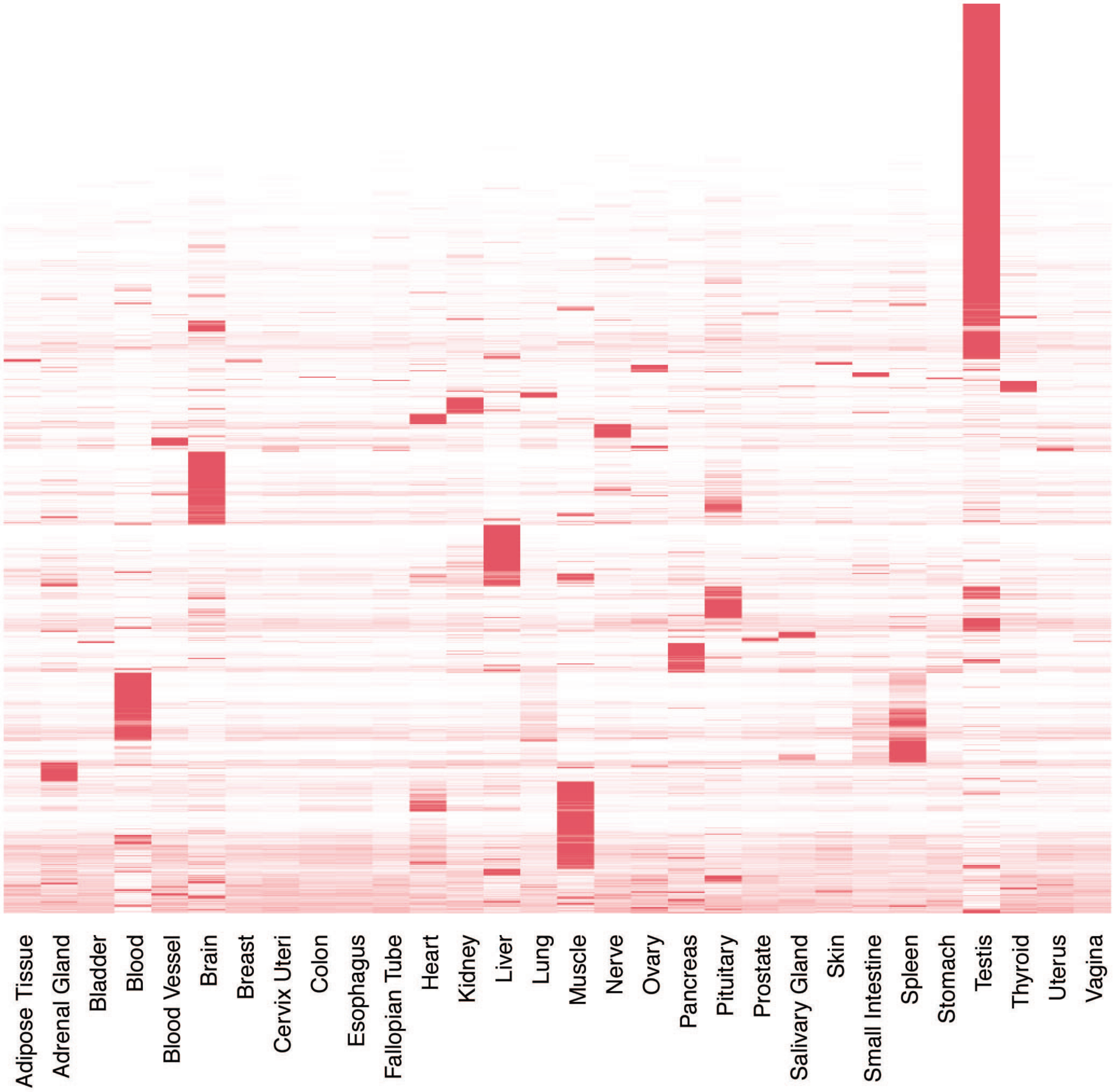


Supplemental Figure 1 Heatmap of the median expression of the specific genes for each tissue, shows the degree of specificity.

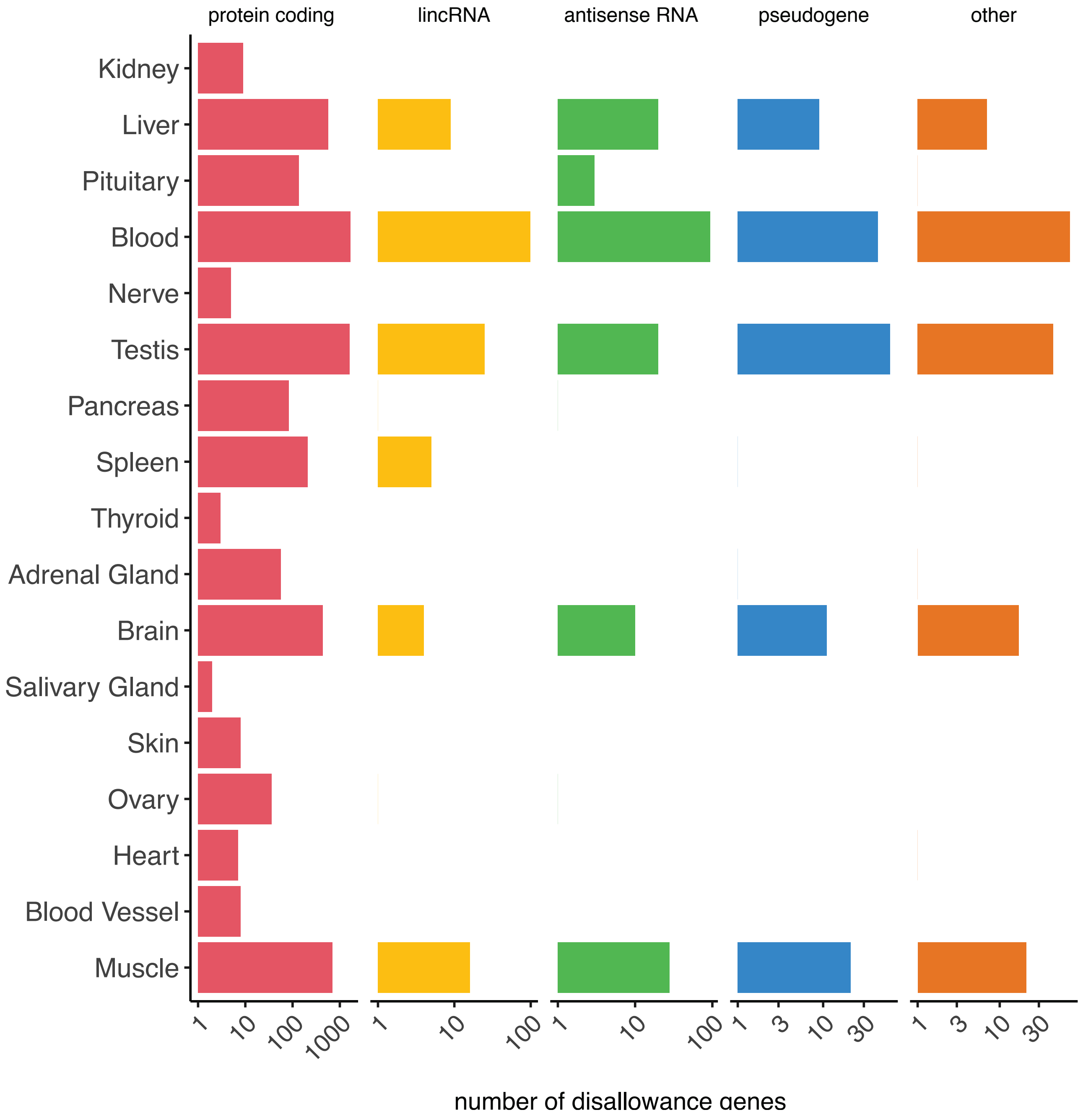


Supplemental Figure 2 Number of disallowance genes for each tissue and biotype is variable.
