## Supplementary material for "SPECS: A non-parameteric method to identify tissue-specific molecular features for unbalanced sample groups": Suplemental Methods

### Empirical Bayes selection of good classifiers based on pairwise comparisons

Olivier Thas

April 8, 2019

#### 1 Introduction

Let the index $d$ refer to a particular sample state (e.g. disease or cancer type), i.e. $d = 1, \ldots, m_d$ with $m_d$ diseases. State $d$ has prevelance $\pi_d$ in the target population. Suppose there are $m_g$ candidate features, i.e. $g = 1, \ldots, m_g$. If no prior knowledge on the prevalences is available, set $\pi_g = 1/m_d$.

Let $Y_{gd}$ denote the outcome of feature $g$ in group $d$. With $n_{gd}$ observations in this group, the individual outcomes are denoted by $Y_{gdi}$, $i = 1, \ldots, n_{gd}$. We use the notation $Y_{g-d}$ to denote the outcome of feature $g$ in all groups but group $d$. When the description of the method applies to a single feature, the index $g$ will be dropped.

A feature is said to be a good classifier or discriminant for a given state, if its outcome distribution for that particular state shows no (or only a little) overlap with the outcome distribution for the other states. This means a large AUC. The AUC is related to the probabilistic index, which is given by

$$p_{gd} = \mathrm{P}\left\{Y_{g-d} < Y_{gd}\right\}.$$

If $p_{gd}$ is very small (close to zero) or very large (close to 1), the outcome distributions in the target state and the other groups are well separated.

The index $g$ will be dropped now. It will be convenient for computational and flexibility reasons to write the probabilistic index as

$$p_d = \mathrm{P}\left\{Y_{-d} < Y_d\right\} = \sum_{k \neq d} \mathrm{P}\left\{Y_k < Y_d\right\} \pi_k.$$

#### 2 Estimation of probabilities

The estimation of probabilities $\mathrm{P}\{Y_k < Y_d\}$ is computationally faster than the estimation of $\mathrm{P}\{Y_{-d} < Y_d\}$. Moreover, if $\mathrm{P}\{Y_{-d} < Y_d\}$ where estimated directly from the sample data, it may be biased because of the state-specific sample sizes may not be proportional to the prevalences in the target population.

In the next few paragraphs we will outline how to estimate the probabilistic indices and how their variances can be estimated consistently. The latter is important for a proper feature selection procedure.

Let $I_{ki;dj} = \mathrm{I}\{Y_{ki} < Y_{dj}\}$ (i.e. a 0/1 indicator). The probability $P_{kd} = \mathrm{P}\{Y_k < Y_d\}$ is then estimated as

$$\hat{P}_{kd} = \frac{1}{n_k n_d} \sum_{i=1}^{n_k} \sum_{j=1}^{n_d} I_{ki;dj}.$$

Hence, an estimator of $p_d = \mathrm{P}\{Y_{-d} < Y_d\}$ is given by

$$\hat{p}_d = \sum_{k \neq d} \hat{P}_{kd} \pi_k.$$

For further purposes we need the sampling distribution of $\hat{p}_d$. We therefore write

$$\hat{\boldsymbol{P}}_d^t = (\hat{P}_{kd})_{k \neq d} \quad \text{and} \quad \boldsymbol{\pi}^t = (\pi_1, \ldots, \pi_{m_d})$$

and

$$\hat{p}_d = \boldsymbol{\pi}^t \hat{\boldsymbol{P}}_d.$$

It's a well known result that each of the $\hat{P}_{kd}$ is asymptotically normal and that they are jointly asymptotically multivariate normal. Hence, $\hat{p}_d$ is asymptotically normal too. Its mean will later be set to 1/2 under the null hypothesis, so at this stage we only need the covariance matrix of $\hat{\boldsymbol{P}}_d$, and a consistent estimator of this covariance matrix.

For $k = l$:

$$\mathrm{Cov}\left\{\hat{P}_{kd}, \hat{P}_{kd}\right\} = \mathrm{Var}\left\{\hat{P}_{kd}\right\}.$$

An estimator of this variance is given in Thas (2010), p. 232-233.

For $k \neq l$:

$$\begin{aligned}
\mathrm{Cov}\left\{\hat{P}_{kd}, \hat{P}_{ld}\right\} &= \frac{1}{n_k n_l n_d^2} \sum_{i=1}^{n_k} \sum_{j=1}^{n_d} \sum_{a=1}^{n_l} \sum_{b=1}^{n_d} \mathrm{Cov}\left\{I_{ki;dj}, I_{la;db}\right\} \\
&= \frac{1}{n_k n_l n_d^2} \sum_{i=1}^{n_k} \sum_{j=1}^{n_d} \sum_{a=1}^{n_l} \mathrm{Cov}\left\{I_{ki;dj}, I_{la;dj}\right\} \\
&= \frac{1}{n_k n_l n_d^2} n_k n_l n_d \mathrm{Cov}\left\{I_{ki;dj}, I_{la;dj}\right\} \\
&= \frac{1}{n_d} \mathrm{Cov}\left\{I_{ki;dj}, I_{la;dj}\right\}
\end{aligned}$$

A further re-expression of $\mathrm{Cov}\left\{I_{ki;dj}, I_{la;dj}\right\}$:

$$\mathrm{Cov}\left\{I_{ki;dj}, I_{la;dj}\right\} = \mathrm{E}\left\{I_{ki;dj} I_{la;dj}\right\} - \mathrm{E}\left\{I_{ki;dj}\right\} \mathrm{E}\left\{I_{la;dj}\right\} = \mathrm{E}\left\{I_{ki;dj} I_{la;dj}\right\} - P_{kd} P_{ld}.$$

Estimates for $P_{kd}$ and $P_{ld}$ are known already. Hence, the for estimation of $\mathrm{Cov}\left\{I_{ki;dj}, I_{la;dj}\right\}$ we only need estimates of $E_{kld} = \mathrm{E}\left\{I_{ki;dj} I_{la;dj}\right\}$. An empirical estimator is given by

$$\begin{aligned}
\hat{E}_{kld} &= \frac{1}{n_k n_l n_d} \sum_{i=1}^{n_k} \sum_{j=1}^{n_d} \sum_{a=1}^{n_l} I_{ki;dj} I_{la;dj} \\
&= \frac{1}{n_k n_l n_d} \sum_{i=1}^{n_k} \sum_{j=1}^{n_d} \sum_{a=1}^{n_l} \mathrm{I}\left\{Y_{ki} < Y_{dj}\right\} \mathrm{I}\left\{Y_{la} < Y_{dj}\right\} \\
&= \frac{1}{n_k n_l n_d} \sum_{i=1}^{n_k} \sum_{j=1}^{n_d} \sum_{a=1}^{n_l} \mathrm{I}\left\{\max(Y_{ki}, Y_{la}) < Y_{dj}\right\}
\end{aligned}$$

With these ingredients, the covariance matrix of $\hat{\boldsymbol{P}}_d$ can be estimated. Let $\hat{\boldsymbol{\Sigma}}$ denote this matrix. An estimate of the variance of $\hat{p}_d$ is then given by

$$\hat{\sigma}_d^2 = \boldsymbol{\pi}^t \hat{\boldsymbol{\Sigma}} \boldsymbol{\pi}.$$

### 3 Selection of good discriminating features

We focus on a single state, say $d$. We reintroduce the index $g$ for feature. Let $\hat{p}_{gd}$ the estimate of $p_{gd}$, and $\hat{\sigma}_{gd}^2$ its variance estimate.

In an empirical Bayesian setting we assume ($g = 1, \ldots, m_g$)

$$p_{gd} \sim g(\cdot)$$

where $g(\cdot)$ is a density function (there will be no need to specify this explicitely).

From the asymptotic normality of $\hat{p}_{gd}$ we can assume

$$\hat{p}_{gd} \mid p_{gd} \sim N(p_{gd}, \sigma^2_{gd}).$$

Let $f_d(\cdot)$ denote the marginal distribution of $\hat{p}_{gd}$ over all features $g = 1, \ldots, m_g$. Tweedie's formula (Efron, 2011) eventually gives

$$\mathrm{E}\left\{p_{gd} \mid (\hat{p}_{gd})_{g=1,\ldots,m_g}\right\} = \hat{p}_{gd} + \sigma^2_{gd} \frac{d}{dz} \log f_d(z)\bigg|_{z=\hat{p}_{gd}}.$$

In this expression the variance $\sigma^2_{gd}$ can be replaced by its estimate, and $f_d(\cdot)$ can be replaced by a nonparametric density estimate upon using all $\hat{p}_{gd}$, $g = 1, \ldots, m_g$. The latter only works with $m_g$ sufficiently large. Let $\tilde{p}_{gd}$ denote this estimate of $\mathrm{E}\left\{p_{gd} \mid (\hat{p}_{gd})_{g=1,\ldots,m_g}\right\}$.

Features can now be selected by ranking the $\tilde{p}_{gd}$ and selecting the largest (closest to 1) or the smallest (closest to zero). The empirical Bayes procedure protects against selection bias (Efron, 2011).
